# Perturbation of GABA Biosynthesis Links Cell Cycle to Control *Arabidopsis thaliana* Leaf Development

**DOI:** 10.1101/2020.02.21.960393

**Authors:** Yaxin Gong, Han Yue, Yu Xiang, Guanghui Yu

**Affiliations:** Hubei Provincial Key Laboratory for Protection and Application of Special Plants in Wuling Area of China, Engineering Research Centre for the Protection and Utilization of Bioresource in Ethnic Area of Southern China, College of Life Sciences, South-Central University for Nationalities, Wuhan 430074, China

**Author notes:** Correspondence: Guanghui Yu.

**Keywords:** *Arabidopsis thaliana*, CDKA;1, endoreplication, GABA, Reactive Oxygen Species

## Abstract

To investigate the molecular mechanism underlying increasing leaf area in γ-Aminobutyric acid (GABA) biosynthetic mutants, the first pair of true leaves of GABA biosynthetic mutants was measured. The results showed that the leaf blade area in GABA biosynthetic mutants was larger than that of the wild type to different extents, and the area of the leaf epidermal cells in mutants was larger than that of the wild type. DNA polyploid analysis showed that polyploid cells in GABA biosynthetic mutants were appearing earlier and more abundant than in the wild type. To check the correlation between cell size and endoreplication, the expression of factors involving endocycles, including D-type cyclin gene (*CYCD3;1, CYCD3;2, CYCD3;3*, and *CYCD4;1*) and kinase *CKDA;1*, were analysed by qRT-PCR. The results showed that *CKDA;1* in GABA biosynthetic mutants was downregulated, and four types of *CYCDs* showed different expression patterns in different GABA biosynthetic mutants. Inconsistent with this result, for *CCS52A* (*CELL CYCLE SWITCH 52A*) (controlling the endocycle entry) in *gad2* and *gad1/gad2* mutants, the expression of *CCS52A2* was significantly higher than that in the wild type. The expression of *SIM* (*SIAMESE*) and *SMR* (*SIAMESE-RELATED*), which inhibit kinase activity, were also upregulated compared with the control. To further study the possible potential relationship between GABA metabolism and endoreplication, we analysed the reactive oxygen species (ROS) levels in guard cells using ROS fluorescent probes. ROS levels were significantly higher in GABA biosynthetic mutants than the control. All results indicated that cyclin, the cyclin*-*dependent kinase, and its inhibitory protein were coordinated to participate in endoreplication control at the transcription level in the leaves of GABA biosynthetic mutant *Arabidopsis*.

**Contribution to the field statement:** γ-Aminobutyric acid (GABA) metabolic pathway plays a dual role in plant development. This research investigated the perturbation of GABA biosynthesis on *Arabidopsis* leave endoreplication for the first time. In the GABA biosynthetic mutants, many genes, participating in cell division regulation, are coordinately transcriptionally expressed to trigger the onset and maintenance of endoreplication, and this led to the cell expansion and the increase leaf blade area. However, this initiation of endoreplication links with the decrease of endogenous GABA level and the increase Reactive oxygen species (ROS). This may be a compensation mechanism to adapt to abnormal GABA level in plant leaf development. Present evidence provided hypothesized that the normal GABA level in plant leaf development plays a brake to inhibit the immature cell expansion and differentiation, and this negative regulation functions a guarantee mechanism to watchdog the normal leaf development. In all, this contribution provides an updated perspective on the role of GABA in plant development.

## 1 Introduction

γ-Aminobutyric acid (GABA) is a four-C, non-protein component amino acid commonly found in organisms and is prevalent in bacteria, plants, and vertebrates (Seifikalhor et al., 2019). The precursor of GABA synthesis is L-glutamic acid (Glu), which is catalysed by glutamate decarboxylase (GAD) in cytoplasm. The *Arabidopsis* genome has five genes encoding GAD named GAD1–5. The oxidative metabolism of GABA occurs in mitochondria and entry is mediated by a GABA permease (Michaeli et al., 2011). In mitochondria, using α-ketoglutarate or pyruvate as an amino acceptor, GABA is catalysed by GABA transaminase (GABA-T) to produce glutamic acid (Glu) or Alanine (Ala) and succinic semialdehyde (SSA), respectively. SSA is oxidized by SSA dehydrogenase (SSADH) to succinic acid (SucA), and then SucA enters the tricarboxylic acid (TCA) cycle for further metabolism. This metabolic pathway is referred to as the GABA shunt. Under hypoxic or high-light conditions, SSA can be reduced to gamma-hydroxybutyrate (GHB) by SSA reductase (SSR, also known as GHB dehydrogenase) in the cytoplasm, mitochondria, and chloroplasts (Allan et al., 2008). Previous studies have reported that unique cytosolic and plastid glyoxylate reductase isoforms in *Arabidopsis* are known as GLYR1 and atGLYR2, respectively, and they catalyse the conversion of SSA to GHB and glyoxylic acid to glycolic acid via an NADPH-dependent reaction (Brikis et al., 2017). The balance of the redox state was maintained by the accumulation of GHB and the reduction of SSA via the GABA shunt.

GABA biosynthesis and accumulation from the glutamic acid pathway and its metabolism are at the junction of N and C metabolism, providing a useful metabolic substrate for the TCA cycle, electron transport chain, and C skeleton, which are involved in the balancing of C:N metabolism (Jacoby et al., 2011). It is generally believed that GABA metabolism is considered to be involved in metabolic signalling, which plays a dual role in plant development, including both metabolic and signal regulation (Häusler et al., 2014; Ramesh et al., 2017; Podlešáková et al., 2019). In addition, aldehyde chemical groups (i.e. H–C=O) produced by GABA biosynthesis and metabolic pathways in plants have high molecular activity, and aldehydes accumulated under stressed conditions are highly toxic and can react with DNA, lipids of oxidative membranes, and modified proteins, or affect the transcription of stress-related genes, leading to cellular and ontogenetic problems in plants. Previous studies have reported that GABA metabolism regulates leaf pattern morphogenesis. The SSADH gene is involved in the formation of the paraxial–abaxial (upper–lower) leaf model in *Arabidopsis thaliana* (Toyokura et al., 2011; 2012). Mutation of *enf1* (*enlarged fil expression domain1*) alters the expression pattern of the *FIL* (*YABBY1, FILAMENTOUS FLOWER*) gene, which is characteristic of meristem and organs in *A. thaliana* (Sawa et al., 1999), on the abaxial surface of leaf primordium (Sawa et al., 1999; Siegfried et al., 1999). However, here is a dearth of research on the role of GABA biosynthesis in the leaf development of *A. thaliana*. In the present study, GABA biosynthetic mutants, *gad1, gad2*, and *gad1/gad2* were examined to explore the molecular mechanism underlying the GABA negative feedback that regulates leaf cell endoreplication during leaf development. Our findings will provide evidence for further understanding the role of GABA in plant development.

## 2 Materials and methods

### 2.1 Experimental Materials

*A. thaliana* Col wild-type seeds, Col ecotype *gad1* mutant, *gad2* mutant, and *gad1/gad2* double mutant were provided by Prof. Barry Shelp (University of Guelph, Canada). The seeds of the wild type and mutants were sterilized for ∼2 h in a sealed container with chlorine gas, and then inoculated on MS solid medium and synchronized for 3 d at 4 °C. After synchronization, the plate was taken out and placed in greenhouse under a photoperiod of 16/8 h light/dark and incubated at 22 °C.

### 2.2 Measurement of leaf blade area

From the 4^th^ day after *A. thaliana* seedlings being transferred to the greenhouse, their growth condition was photographed every 24 h with a Brinno time-lapse camera (TLC100) at a fixed distance to record the leaf blade area.

### 2.3 Microscopic observation and measurement of cell area

From the 7^th^ day after *A. thaliana* seedlings being transferred to the greenhouse, the first pair of true leaves was collected daily, the leaf abaxial epidermis located 25% and 75% from the distance between the tip and the base of the leaf blade was photocopied with nail polish, and then photographed using an Olympus DP80 microscope, and the area was calculated with the software that came with the microscope. Under the microscope, a certain area of the epidermis was confined to count the number of cells, and then the average cell area was calculated.

### 2.4 DNA ploidy analysis

Approximately two or three leaves at the young stage or one or two leaves at the middle and late leaf developmental stages were chopped with a razor blade in nuclei extraction buffer (CyStain^®^ UV Precise P, Sysmex Partec), and then transferred to the staining buffer (CyStain^®^ UV Precise P, Sysmex Partec) according to the manufacturer’s instructions. The ploidy level of DNA in leaf cells was determined using a CyFlow Ploidy Analyser (Sysmex Partec).

### 2.5 qRT-PCR analysis

Total RNA from *A. thaliana* leaves was extracted by AxyPrep Multisource Total RNA Miniprep Kit (Axygen Science, Inc). Reverse transcription was performed by Goldenstar RT6 Gene Synthesis Kit (Tingke Biotechnology Co., Ltd.) with a reverse transcription reaction system of 20 μL. qRT-PCR analysis was performed by MyGo Pro qPCR System (IT-IS Life Science Ltd.). The gene *isoprenyl diphosphate delta isomerase II* (*IPP2*, AT3G02780) was selected as the internal reference gene according to previous research (Fung-Uceda et al., 2018). The primers of the genes involved in cell cycle regulation, endocycle initiation, progression, and exit are listed in the supporting materials (Table S1).

### 2.6 *In vivo* reactive oxygen species imaging

For GC reactive oxygen species (ROS) staining, H2DCF-DA stock solution (10 mmol/L stock in Dimethyl Sulfoxide) was diluted in deionized water to yield a final concentration of 6.25 μmol/L with a final dimethyl sulfoxide concentration of 0.0125% (v/v) (Watkins et al., 2017). Epidermal strips were peeled with adhesive tape and directly stained for 30 min in the above solution. After rinsing with deionized water for 5 min, images were taken by Olympus DP80 fluorescent microscopy. The intensity of coloration was quantified using ImageJ software (National Institutes of Health, USA).

### 2.7 Statistical analyses

Data are expressed as averages ± standard error (SE). Experiments were conducted with two or three independent replicates. One-way ANOVA was employed using SPSS 20.0.

## 3 Results

### 3.1 The leaf blade area of the GABA biosynthetic mutant was larger than that of the wild type

At the early stage of leaf development (4^th^ day after stratification), there was no significant difference in the area of the first pair of true leaves between the wild-type and GABA biosynthetic mutants (Figure 1). As time progressed, the area of the mutant leaves increased more rapidly than that of the wild type. Around the 8^th^ day, the leaf area of the first pair of true leaves in the wild type was about 1.8 mm^2^, and the first pair of true leaves in *gad1* and *gad2* mutants was about 2.4 mm^2^, at 1.3-fold that of the wild type, and that of the *gad1/gad2* mutants was about 3.2 mm^2^ with 1.8-fold that of the wild type (Figure 1). The leaf area between the GABA biosynthetic mutants and wild type was significantly different (*P* <0.05).

**Figure 1.**
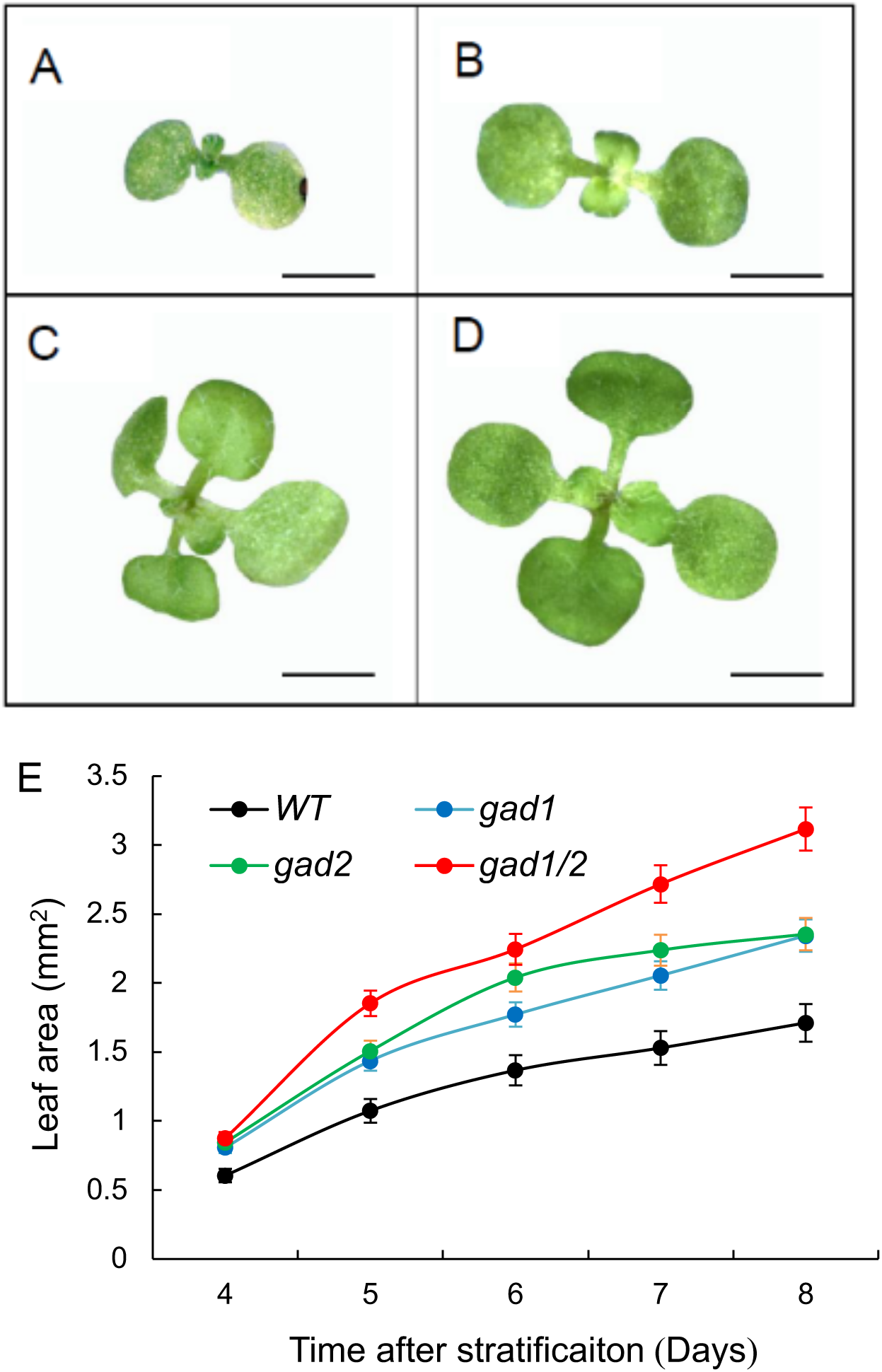
Leaf area of γ-Aminobutyric acid (GABA) biosynthetic mutants was larger than that of wild-type Arabidopsis. Top view of leaves at 8 d after stratification of A. wild type; B. *gad1* mutant, C. *gad2* mutant and D. *gad1/gad2* mutants; E. Leaf growth curve of GABA mutants and wild type of Arabidopsis in the early stage (from 4–8 d after stratification). Results are presented as averages ± SE of three separate experiments (n = 15). Bar in A–D = 860 μm.

### 3.2 Cell size in GABA biosynthetic mutants was larger than those of the wild type

To determine the reasons for the leaf area in GABA biosynthetic mutants being larger than that in the wild type, we compared the area of leaf epidermal cells. The average cell area in the epidermis of the first pair of true leaves of the wild type was about 2700 μm^2^ at the 8^th^ day after stratification (Figure 2). The average cell area of the epidermis of the first pair of leaves of *gad1* and *gad2* mutants was ∼3200 μm^2^, 1.2-fold that of the wild type (*P* <0.05). The average cell area of the epidermis of the first pair of true leaves in *gad1/gad2* mutants was ∼3800 μm^2^, and was 1.4-fold that of the wild type with significant differences (*P* <0.05) (Figure 2).

**Figure 2.**
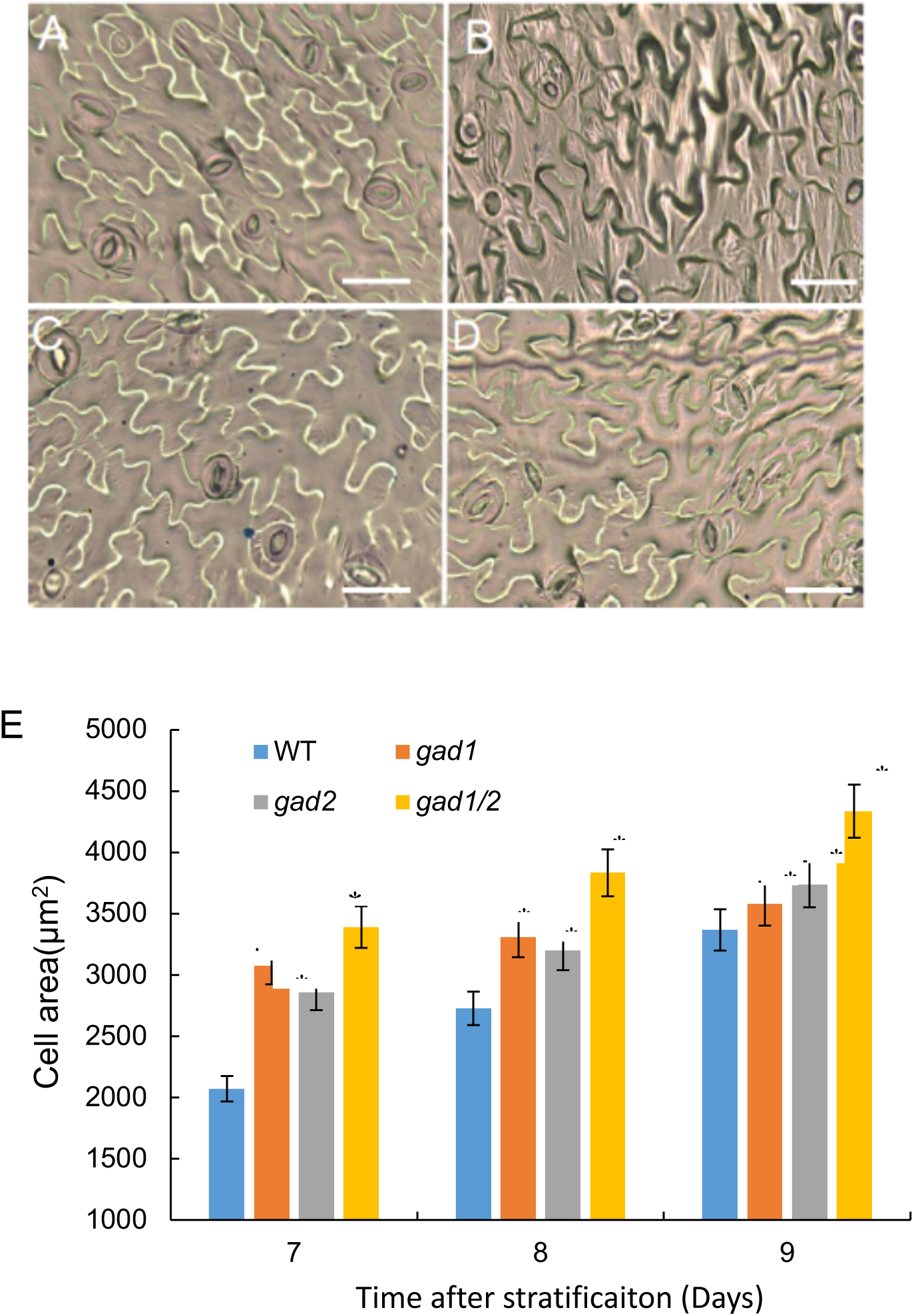
Epidermal cell size in the first true leaves of γ-Aminobutyric acid (GABA) biosynthetic mutants was larger than that of wild-type Arabidopsis at the early growth stage. Epidermal cell of A. wild type; B. *gad1* mutant, C. *gad2* mutant and D. *gad1/gad2* mutants; E. Cell size comparison among GABA mutants and wild type of Arabidopsis in the early stage (7–9 d after stratification). Results are presented as averages ± SE of 60 cells calculated across three separate experimental replicates. Asterisks represent significant differences between the mutants and the wild type (*P* <0.05) and were determined using one-way ANOVA. Bar in A–D = 55 μm.

### 3.3 The polyploidy in GABA biosynthetic mutant leaf occurred earlier and higher than that of the wild type

The correlation between cell size and DNA ploidy level (Matsunaga et al., 2013) prompted us to explore whether the cell size of GABA biosynthetic mutants is related to DNA polyploidy level. On the 7^th^ day of leaf development after stratification, the cells in the first pair of true leaves in wild-type *A. thaliana* were mostly diploid and tetraploid, indicating that leaf cells mainly divided at this stage (Figure 3A). However, leaf cells in *gad1, gad2*, and *gad1/gad2* mutants appeared to be in different proportions of 8-ploid cells except the diploid and tetraploid cells (Figure 3B). The appearance of 8-ploid cells in leaves is a marker of endoreplication. On the 8^th^ day of leaf development, 8-ploid cells were not found in wild-type *A. thaliana*, and the proportion of octoploid cells in three kind of GABA-biosynthetic mutants increased further (Figure 3B).

**Figure 3.**
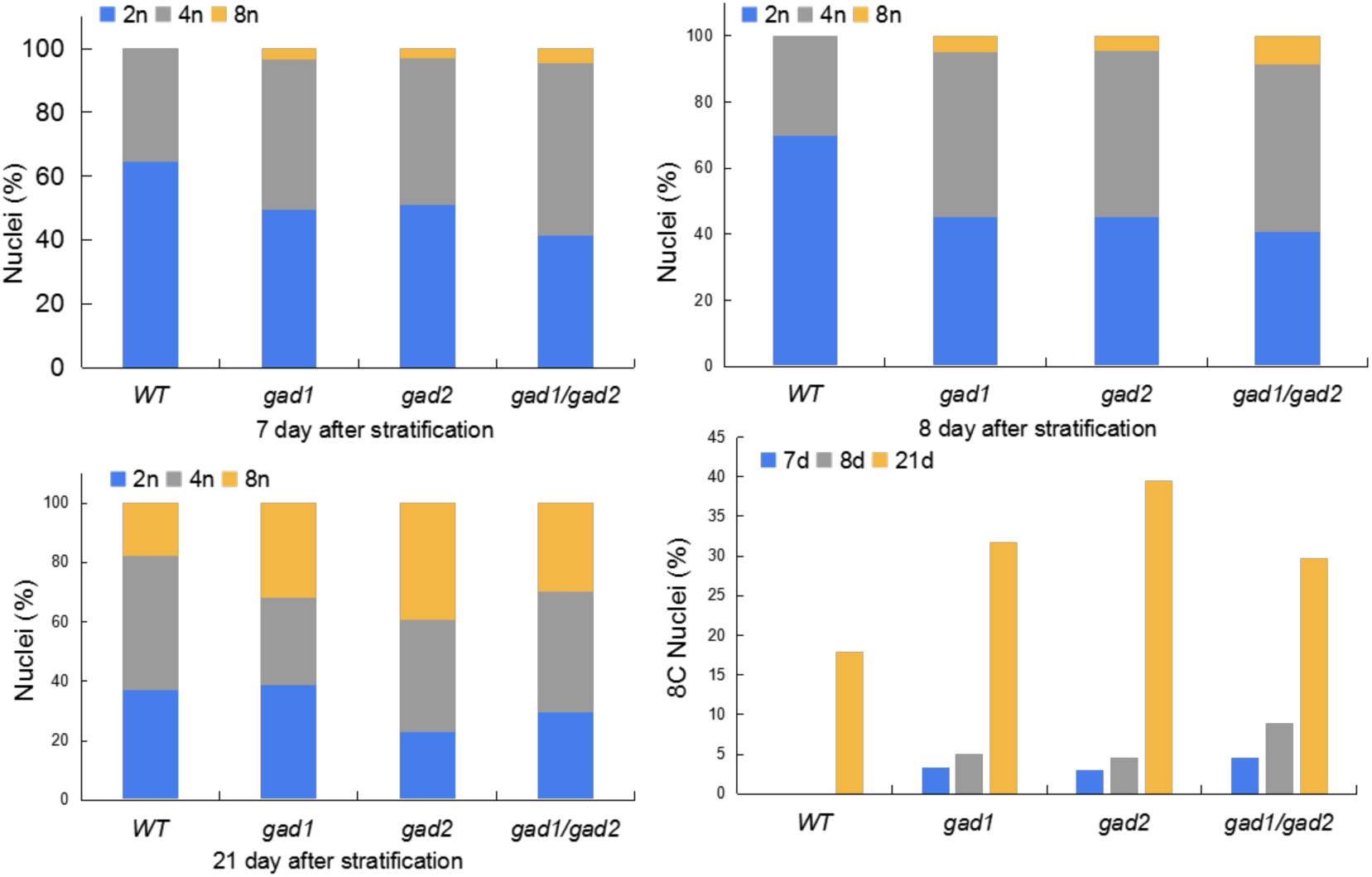
Octoploid (8n) cells occurred earlier in biosynthetic mutants than that in the wild type. Top panel: DNA ploidy analysis in leaf early growth stage (7–8 days after stratification) between γ-Aminobutyric acid (GABA) biosynthetic mutants and the wild type. Left bottom panel: DNA ploidy analysis in leaf middle-late stage (21 days after stratification) between GABA mutants and the control. Right bottom panel: 8C percentage comparison among mutants and the wild type during the long growth period.

To further confirm the relationship between cell enlargement and cell ploidy level, we compared the cell ploidy level in the middle and late leaf development stages (21^st^ day after stratification). The proportion of octoploid cells in the wild type was about 18% (Figure 3C). However, the proportion of the octoploid cells in *gad1, gad2*, and *gad1/gad2* mutants was about 30–40%, which was significantly higher than that in the wild type (Figure 3C).

Overall, at the early stage of leaf development (7–8^th^ day after stratification), the 8-ploid cells were only detected in the GABA biosynthetic mutants (8-ploid cells in *gad1* and *gad2* mutants was 3–5%, and that in *gad1/gad2* double mutants was about 5–9%) (Figure 3D). At the late stage of leaf development (21^st^ day after stratification), the proportion of 8-ploid cells in wild-type leaves was also detected; however, its level was significantly lower than that in *gad* mutants (Figure 3D).

### 3.4 Type-D cyclin genes involved in endocycle regulation in GABA biosynthetic mutants

Considering that endoreplication is a special form of cell division (De Veylder et al., 2011) and is the reason for cell polyploidy (Matsunaga et al., 2013), type-D relative gene participation in cell cycle regulation was analysed by qRT-PCR.

CYCD3;1 (At4g34160) is a key component to balance cell proliferation/division and endoreplication (Dewitte et al., 2007). Downregulation of *CYCD3;1* can lead to endoreplication, and its overexpression can cause excessive cell proliferation and inhibit cell differentiation (Dewitte et al., 2003). qRT-PCR confirmed that the expression of *CYCD3;1* was significantly downregulated in GABA biosynthetic mutants (Figure 4). In *gad1* and *gad2* single mutants, the relative expression of *CYCD3;1* was about 0.2-fold that of the wild type (*P* <0.01), and in *gad1/gad2* double mutants, the relative expression of *CYCD3;1* was about 0.1-fold that of the wild type (*P* <0.01) (Figure 4), indicating that blocking GABA synthesis inhibited the expression of *CYCD3;1*.

**Figure 4.**
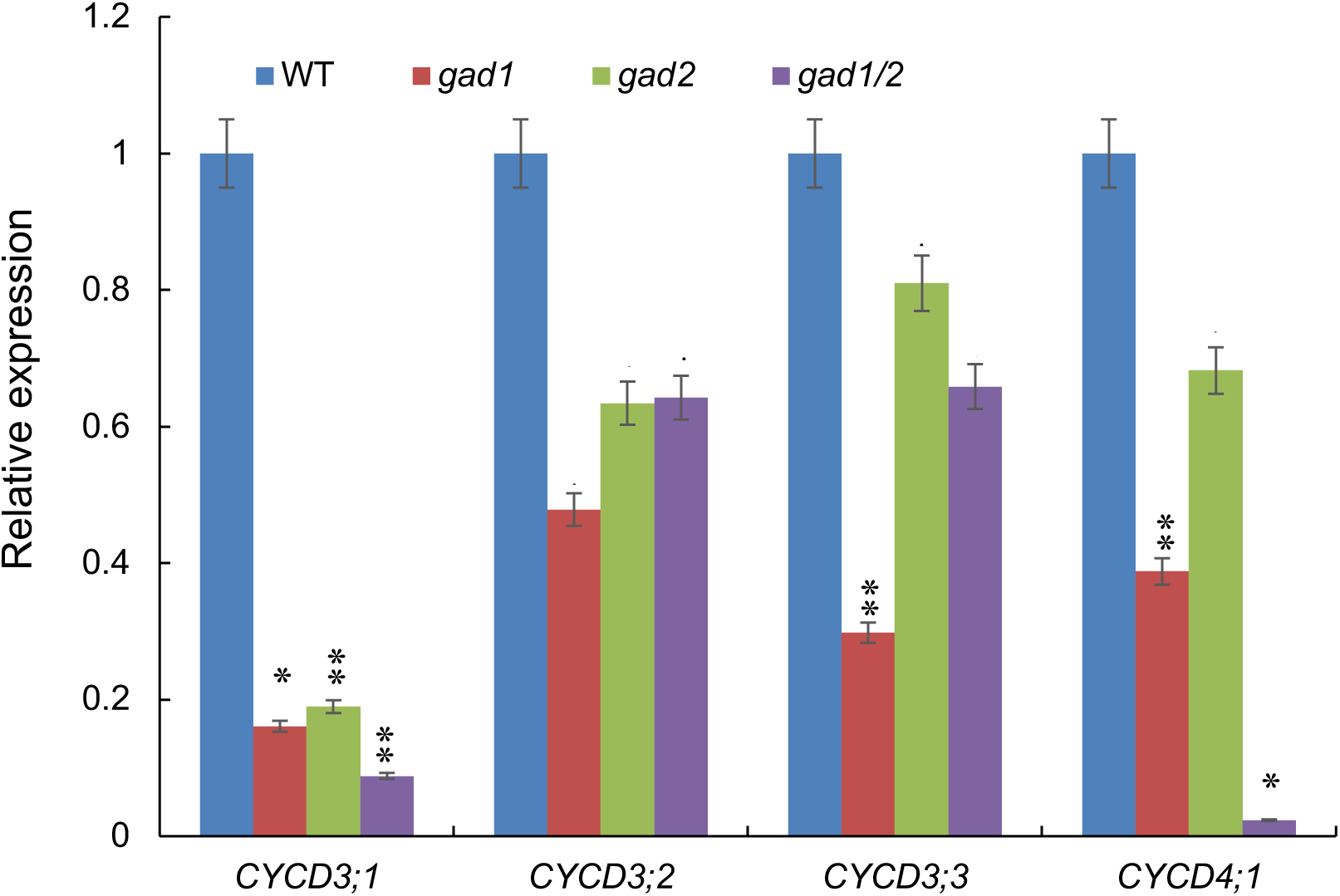
Expression of *Cyclin D* components in biosynthetic mutants was lower than that in the wild type at the 8^th^ day after stratification. Results are presented as averages ± SE of triplicate experiments. Asterisks represent significant differences between the mutants and the wild type (\**P* <0.05; ** *P* <0.01) and were determined using one-way ANOVA.

CYCD3;2 (At5g67260) and CYCD3;3 (At3g50070) participate in the symmetrical division of guard mother cells and guard cells (GCs) in the late stomatal lineage development (Yang et al., 2014). In the *gad1* mutant, the expression of *CYCD3;2* was significantly downregulated (*P* <0.05); in the *gad2* and *gad1/gad2* mutants, the relative expression of *CYCD3;2* was about 0.6-fold that of the wild type with significant differences (*P* <0.01) (Figure 4). Similarly, the expression of *CYCD3;3* in the *gad1* mutant was about 0.3-fold that of the wild type (*P* <0.01), and in the *gad2* and *gad1/gad2* mutants, the expression of *CYCD3;3* was 0.7 and 0.5-fold that of the wild type, respectively (*P* <0.05) (Figure 4). CYCD4;1 (At5g65420) could activate the cell cycle of root apical meristem (Masubelele et al., 2005), and the expression of CYCD*4;1* in the *gad2* mutant was significantly downregulated (*P* <0.05), and in the *gad1* and *gad1/gad2* mutants the relative expression of *CYCD4;1* decreased to become the most significantly different to that of the wild type (*P* <0.01) (Figure 4).

### 3.5 The expression of *CDKA;1* in GABA biosynthetic mutants was significantly downregulated

CDKA;1 (At3g48750) is one of the core components of cell cycle regulation, mainly controlling the transition of the G1/S and G2/M mitotic phase (Nowack et al., 2012). Inhibition of its activity can block mitosis to initiate leaf cell endoreplication (Verkest et al., 2005). The expression of *CDKA;1* in GABA biosynthetic mutants was significantly lower than that in the wild type (*P* <0.05). These results indicated that blocking GABA synthesis is linked with the decreased expression of *CDKA;1* (Figure 5).

**Figure 5.**
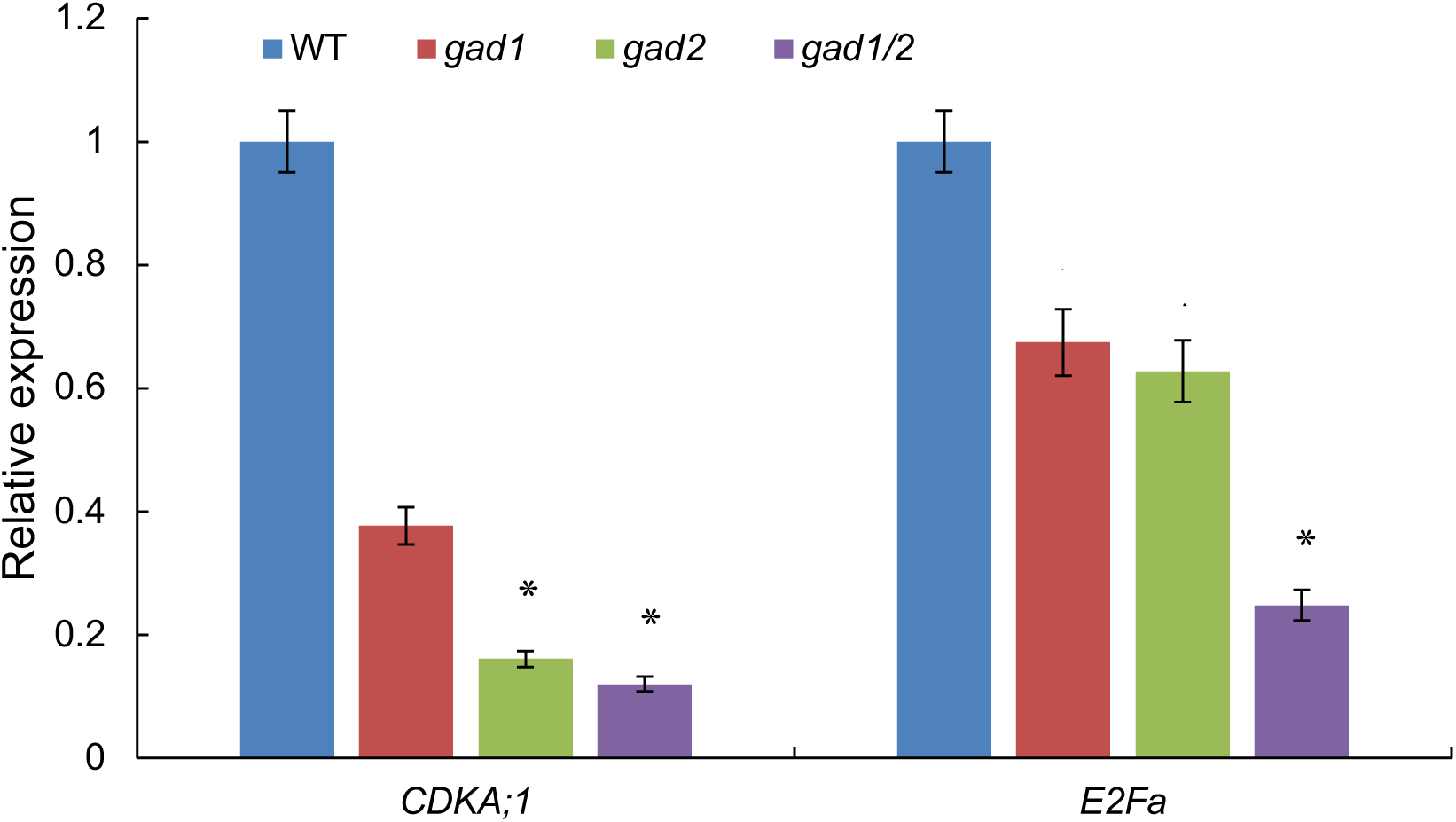
Expression of *CDA;1* and *E2Fa* in biosynthetic mutants was lower than that in the wild type at the 8^th^ day after stratification. Results are presented as averages ± SE of three independent experimental replicates. Asterisks represent significant differences between the mutants and the wild type (\**P* <0.05; ** *P* <0.01) and were determined using one-way ANOVA.

The transcription factor E2Fa (At2g36010) stimulates cell proliferation and delayed differentiation (Boudolf et al., 2004). In the *gad1* and *gad2* mutants, the expression of *E2Fa* was significantly downregulated (*P* <0.05). In *gad1*/*gad2*, the level of *E2Fa* was 0.2-fold that in the wild type (*P* <0.01) (Figure 5).

### 3.6 Different expression patterns of *CCS52A2* and *CDC6* were observed in GABA synthetic mutants

*CCS52A2* (*CELL CYCLE SWITCH52*, At4g11920) encodes a ubiquitin ligase regulating cell cycle division phase (M) and a substrate-specific activator of anaphase promotion complex/cyclosome (Fülöp et al., 2005). Inhibition of kinase activity after *CCS52A2* expression and enhancement of its activity are prerequisites for endoreplication (Heyman et al., 2017; Umeda et al., 2019). Mutations in the *CCS52* gene resulted in delayed endoreplication, and its overexpression resulted in increased DNA ploidy levels (Heyman et al., 2017). In the *gad1* mutant, the relative expression of *CCS52A2* was not different from that of the wild type; however, in the *gad2* and *gad1/gad2* mutants, the expression of *CCS52A2* was significantly higher than that of the wild type (*P* <0.05) (Figure 6).

**Figure 6.**
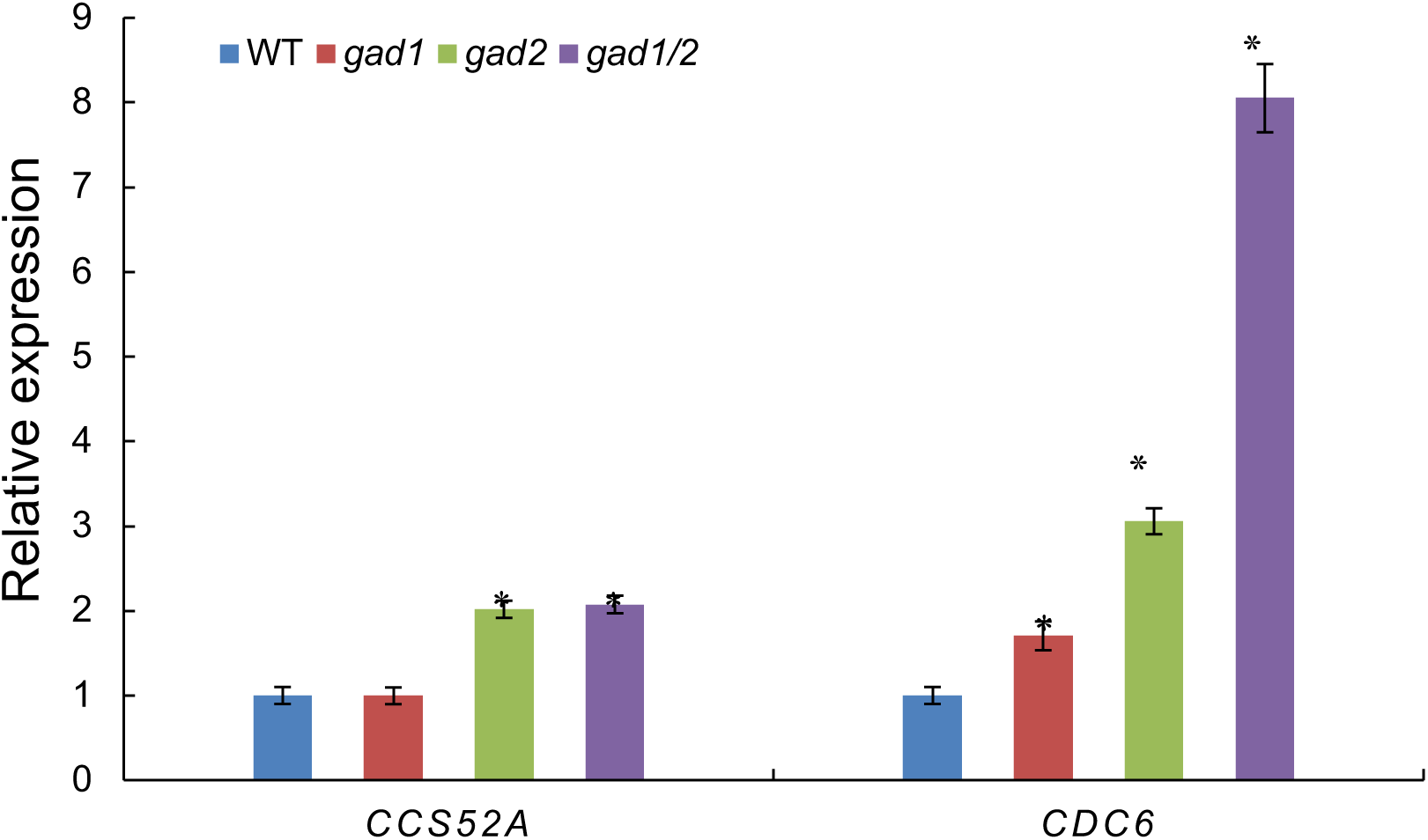
Expression of *CCS52A* and *CDC6* in biosynthetic mutants was higher than that in the wild type at the 8^th^ day after stratification. Results are presented as averages ± SE of three independent experimental replicates. Asterisks represent significant differences between the mutants and the wild type (\**P* <0.05; ** *P* <0.01) and were determined using one-way ANOVA.

*CDC6* (At2g29680) encodes a homolog of cell division regulatory protein 6 (Castellano et al., 2001), which is a license gene for DNA replication (Fung-Uceda et al., 2018). CDC6 participates in the initiation of DNA replication and is an important factor for maintaining endoreplication. The ectopic expression of *CDC6* can increase the ploidy level caused by endoreplication (Castellano et al., 2001; 2004). In the *gad1* mutant, the relative expression of *CDC6* was 1.7-fold as much as that in the wild type (*P* <0.05) (Figure 6). The expression of *CDC6* in the *gad2* mutant was about 3-fold that in the wild type (*P* <0.01), and the expression of *CDC6* in the *gad1/gad2* double mutant was about 8-fold that in the wild type (*P* <0.01) (Figure 6).

### 3.7 The expression of *SIAMESE* and *SIAMESE-RELATED* in GABA biosynthetic mutants was significantly up-regulated

SIAMESE (SIM, At5g04470) is an inhibitory protein of cyclin-dependent kinase and a regulator of mitotic inhibition and endoreplication (Churchman et al., 2006). Overexpression of *SIM* results in dwarfing of plants, serrated leaves, and cells with higher nuclear DNA content (Churchman et al., 2006). The relative expression of the *SIM* in *gad1* single mutant is about twice that of the wild type (*P* <0.05), in the *gad2* single mutant is about 4-fold that of the wild type (*P* <0.05), and in the *gad1/gad2* double mutant is about 7-fold that of the wild type (*P* <0.01) (Figure 7).

**Figure 7.**
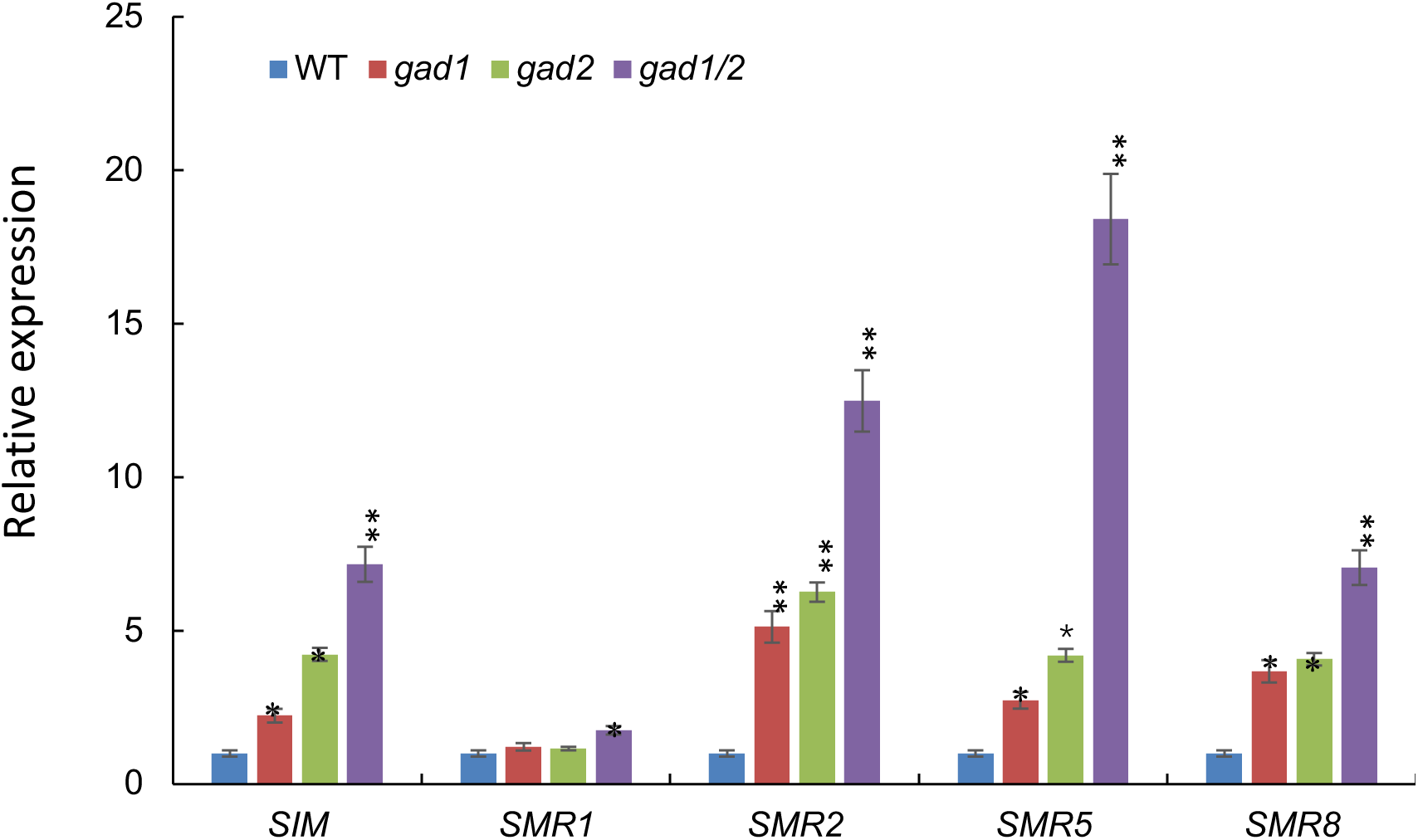
Relative expression of *SIM* gene. Expression of *SIAMESE* (*SIM*) and *SIAMESE-RELATED* (*SMR*) components in biosynthetic mutants was higher than that in the wild type at the 8^th^ day after stratification. Results are presented as averages ± SE of three independent experimental replicates. Asterisks represent significant differences between the mutants and the wild type (\**P* <0.05; ** *P* <0.01). Statistics were determined using one-way ANOVA.

Most components of the *SIAMESE-RELATED* (*SMR*) gene family function in mitosis inhibition and endocycle promotion (Yi et al., 2014; Kumar et al., 2015; Dubois et al., 2018). In the chosen genes, the expression of *SMR1* (At3g10525) in *gad1* and *gad2* mutants was not different to that of the wild type. In the *gad1/gad2* mutants, its expression is 1.5-fold that of the wild type (*P* <0.05) (Figure 7). However, the expression of other components, *SMR2* (At1g08180), *SMR5* (At1g07500), and *SMR8*, was markedly different to that of the wild type (Figure 7). The expression of *SMR2* in *gad1, gad2*, and *gad1/2* reached 5-, 6-, and 11-fold of that in the wild type, respectively (*P* <0.01). Similarly, the expression of *SMR5* in *gad1, gad2*, and *gad1/2* reached 3-, 5- (*P* <0.05), and 17-fold that in the wild type (*P* <0.01), respectively, and the expression of *SMR8* in *gad1, gad2*, and *gad1/2* reached 4-, 4- (*P* <0.05), and 6-fold that in the wild type (*P* <0.01), respectively.

### 3.8 The level of ROS in leaves of GABA biosynthetic mutant was higher than that of the wild type

The fact that redox regulates cell proliferation and the cell cycle (Schippers et al., 2016), and GABA could scavenge ROS (Liu et al., 2011; Seifikalhor et al., 2019), led us to postulate that ROS levels in GABA biosynthetic mutants could be increased. Imaging of H2DCFDA (DCFH-DA; 2’, 7’-Dichlorodihydrofluorescein diacetate) with the fluorescent probe of ROS showed that the average ROS level intensity in GCs of *gad* mutants was higher than that in the wild type (*P* <0.05) (Figure 8). These results suggest that GABA synthetic mutants are indeed correlated with ROS accumulation.

**Figure 8.**
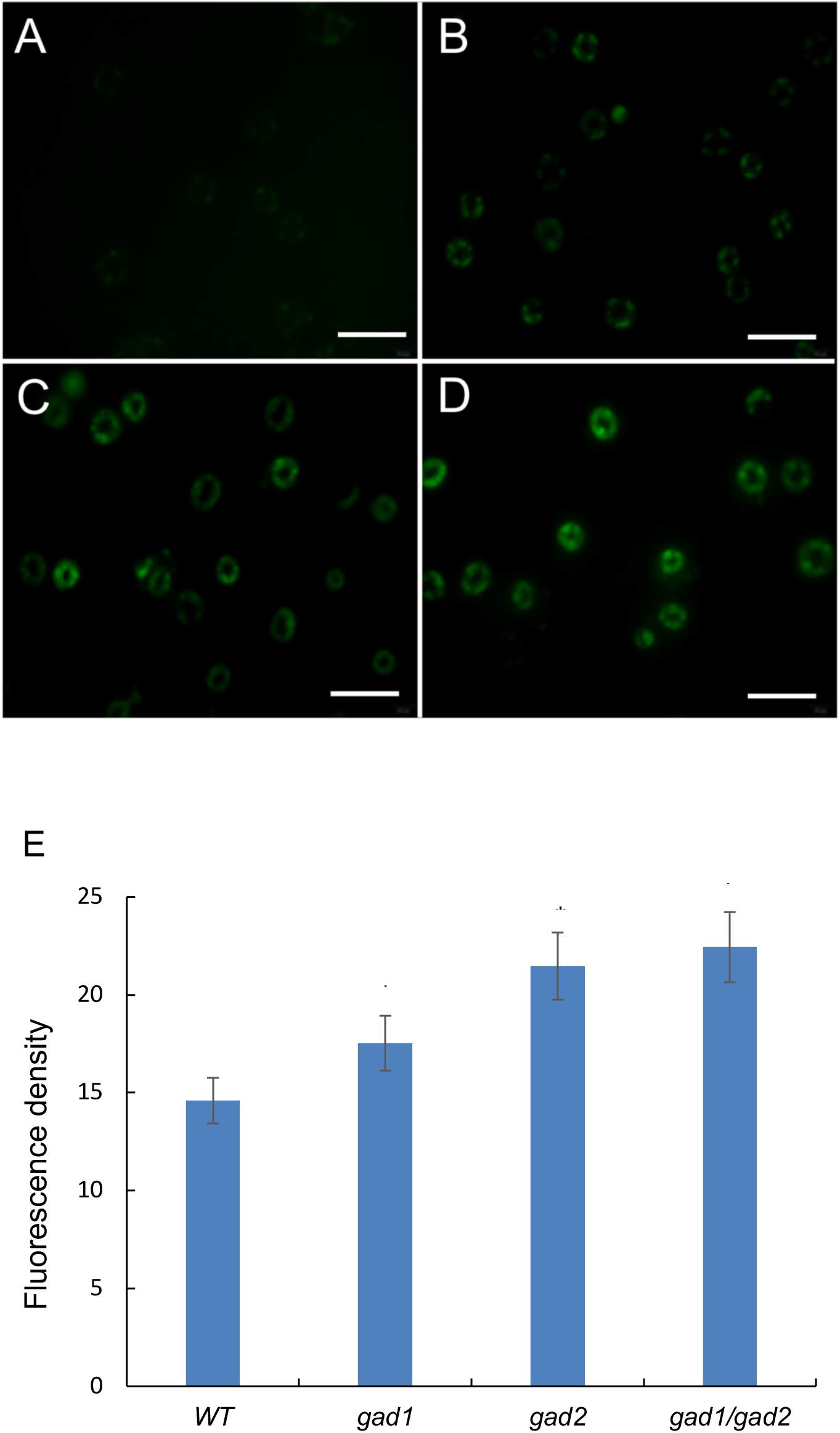
Reactive oxygen species (ROS) level in guard cells (GCs) of γ-Aminobutyric acid (GABA) mutants are higher than that of control A–D. DCF fluorescence (green) in epidermal GCs of GABA biosynthetic mutants and wild type Arabidopsis. E. DCF fluorescence was quantified in the mutants and control. Results are presented as averages ± SE of three separate experiments (n = 16). Asterisks represent significant differences between the mutant and the wild type (*P* <0.05) and were determined using one-way ANOVA. Bar in A–D = 55 μm.

## 4 Discussion

The GABA metabolic pathway plays a dual role during plant development, including metabolic and signal regulation (Bown & Shelp, 2016; Seifikalhor et al., 2019). Abnormal GABA biosynthesis and metabolism have important effects on plant development (Baum et al., 1996; Palanivelu et al., 2003; Renault et al., 2013). The expression of the *Petunia* GAD gene, which lacked the calmodulin-binding domain in transgenic tobacco, resulted in abnormal plant development, which was shorter and more branched than that of normal plants (Baum et al., 1996). In the Arabidopsis *GABA-T* (*pop2-1*) mutant, the growth of pollen tubes could not accurately target the ovule sac in the pistils (Palanivelu et al., 2003). The molecular mechanism was related to the destruction of the gradient distribution of GABA in the stigma of the pistils (Palanivelu et al., 2003). Furthermore, the growth retardation in hypocotyl epidermal cells and root cortex cells is related to the limitation of cell elongation (Renault et al., 2011), and inadequate expression of *GABA-T* may lead to developmental defects in roots and hypocotyls and composition change in cell walls (Renault et al., 2013).

In the present study, the molecular mechanism underlying leaf-area enlargement in GABA biosynthetic mutants was investigated. The growth of plant leaves is largely limited by the development of the epidermis. Polyploidy in pavement cells is strongly correlated with cell size (*P* <0.01) (Melaragno et al., 1993). We confirmed that the larger leaf area and cell size in GABA biosynthetic mutants (Figures 1, 2) was related to much higher percentages of polyploid cells in GABA biosynthetic mutant leaves, which occurred earlier and in higher abundance than that of the wild-type leaf cells (Figure 3).

Endoreplication is a special kind of cell division (De Veylder et al., 2011), and the essence of endoreplication is to maintain CDK activity below the threshold that triggers mitosis (De Veylder et al., 2011). During leaf development of *A. thaliana*, the decrease in transcription levels of mitotic CDK and cyclin genes resulted in endoreplication (Beemster et al., 2006). Consistent with these results, the qRT-PCR analysis confirmed that the type-D cyclin genes (*CYCD3;1, CYCD3;2, CYCD3;3*, and *CYCD4;1*) were differently downregulated in GABA biosynthetic mutants, and the expression of *CDKA;1* exhibited a decreasing expression trend (Figures 4, 5).

Other factors interrelated with endoreplication regulation, from endoreplication initiation, progression, and maintenance, and exit (Breuer et al., 2014), synergistically regulate the occurrence of endoreplication in the leaves of GABA biosynthetic mutants. *CCS52A* (*CELL CYCLE SWITCH 52A*) plays an important role in cell cycle exit and endoreplication entry (Lammens et al., 2009; Vlieghe et al., 2005). The concentration of *CCS52A* in mitotic cells remains below a critical threshold to prevent immaturely initiating endoreplication (Lammens et al., 2009; Vlieghe et al., 2005). In the *gad2* and *gad1/gad2* mutants, the relative expression of *CCS52A2* was significantly higher than that of the wild type (Figure 6). The expression of cyclin-dependent kinase inhibitor *SIM* (*SIAMESE*) and its *SMR* (*SIAMESE-RELATED*) was significantly higher than that of the control (Figure 7). These results demonstrated that the endoreplication regulators, including cyclin, cyclin-dependent kinase, and kinase inhibitor, at the transcriptional level, all harmoniously contributed to the regulation of endoreplication in mutants.

It is worth noting that *SMR5*, a member of *SMR* (*SIAMESE-RELATED*), is an ROS-induced gene, which is more highly expressed in GABA biosynthetic mutants (3–17-fold that of the control) (Figure 7). ROS is an important signalling molecule regulating leaf development and is involved in triggering endoreplication (Schippers et al., 2016). Previous evidence proved that exogenous GABA could scavenge ROS (Liu et al., 2011), and the GABA level in mutants was much lower than that in the wild type (Figure S1); thus, we speculated that the perturbation of GABA biosynthesis may be intrinsically linked with the ROS level. Detection using an ROS fluorescence probe confirmed that the level of ROS in GCs of GABA biosynthesis mutants was significantly higher than that in the control (Figure 8). Therefore, we speculated that blocking the GABA metabolic shunt pathway will lead to the accumulation of ROS intermediates (Bouché et al., 2003; Fait et al., 2005), which may trigger endoreplication (Figure 9). In this hypothesis, normal GABA metabolism functioned as a signal and antioxidant to effectively inhibit the production of ROS. In the GABA biosynthetic mutants, this inhibition was relieved. The production of ROS promotes the expression of *SIAMESE* (*SIM*) and *SIAMESE*-*RELATED* (*SMR*) expression, of which, their encoding products inhibited kinase activity to initiate endoreplication (Figure 9). In contrast, normal GABA metabolism postponed endoreplication, and this negative regulation functions to prevent premature cell differentiation and to ensure normal leaf development. However, this conclusion must be made cautiously because, based on current data, it is difficult to demonstrate a causal relationship between GABA level and ROS in terms of triggering endoreplication during the development of Arabidopsis leaves.

**Figure 9.**
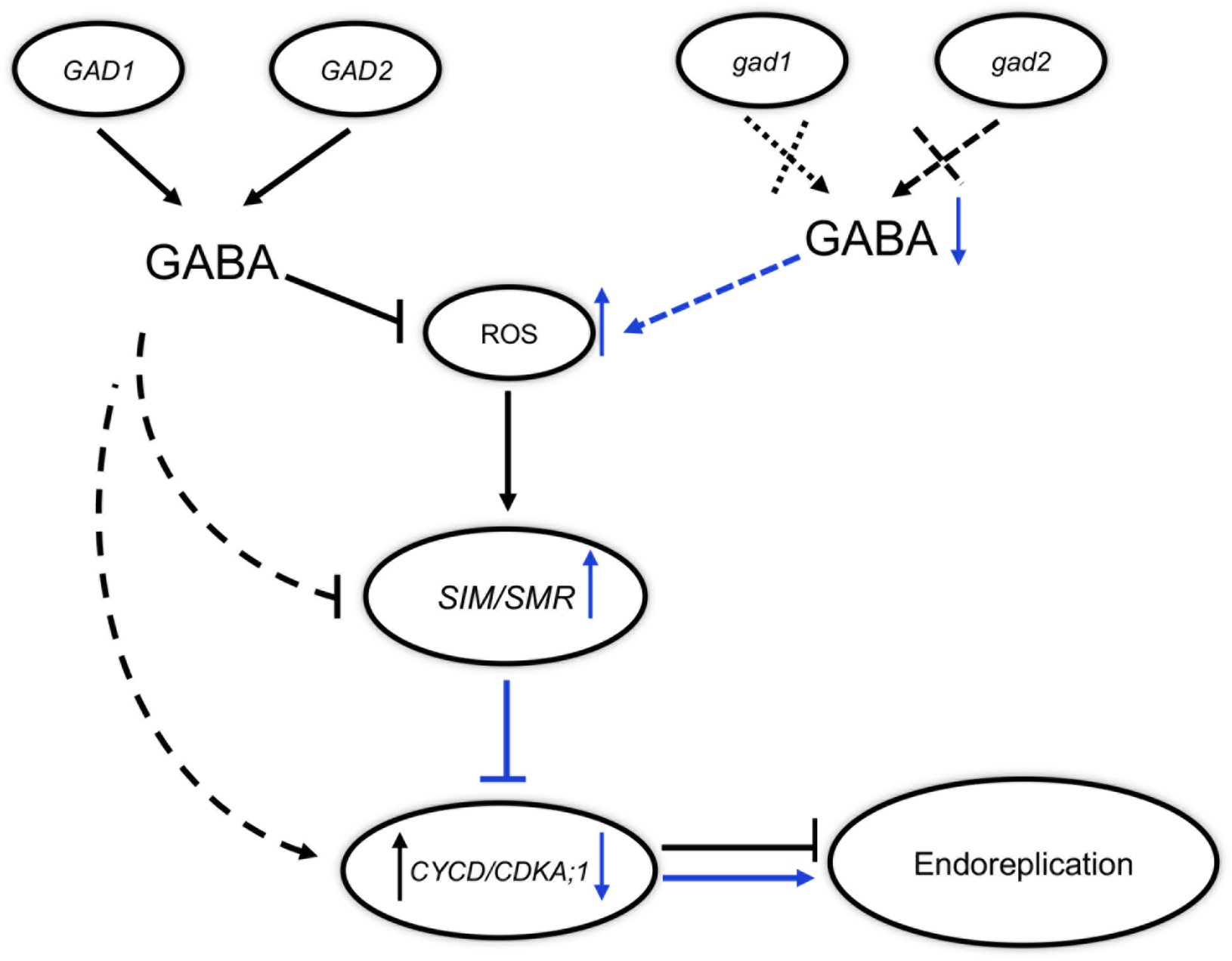
Working hypothesis of γ-Aminobutyric acid (GABA) negatively control the endoreplication of Arabidopsis leaves. GABA, as a normal metabolic signal and antioxidant, effectively inhibits reactive oxygen species (ROS) production. In the GABA biosynthetic mutants, owing to biosynthetic perturbation, the inhibition of GABA on ROS was relieved. The production of ROS promotes the expression of *SIAMESE* (*SIM*) and *SIAMESE*-*RELATED* (*SMR*). Both components are inhibitors of Cyclin-dependent Kinase complexes (e.g. CYC3;1-CDKA;1), and low levels of kinases initiate endoreplication in Arabidopsis. Other components, such as CDC6, synergistically respond to low levels of GABA to prime endoreplication. In the wild type, the normal metabolism of GABA delays endoreplication and prevents premature endoreplication.

Overall, the results of the present study demonstrate that the increase in polyploidy level in GABA biosynthetic mutant leaves is achieved through endoreplication, which is reflected in the increase in average cell size (Breuer et al., 2010). In this regulation, multiple regulators are involved in the initiation, maintenance, and exit of endoreplication. However, the premature termination of mitosis and the immature occurrence of endoreplication are perhaps a compensatory mechanism for leaf immature development, which resulted in increased cell ploidy. This mechanism may be mediated by ROS signalling, but the complex regulatory mechanism requires further research.

## 5 Conclusion

Using GABA biosynthetic mutants, the present study focused on the perturbation of GABA biosynthesis on endoreplication in Arabidopsis leaf development. The endoreplication cells that occurred in mutants were earlier and of higher abundance than those in the wild type. This is the reason to lead to the increase in cell size and leaf blade area. For transcription-level regulation, qRT-PCR confirmed that many genes involved in cell cycle regulation were synergistically participating in the initiation, progression, and maintenance of endoreplication. Among the regulators, *SMR5*, encoding protein inhibited CDK activity, was markedly upregulated in GABA biosynthetic mutants. Owing to the perturbation of GABA biosynthesis, the content of ROS increased in GABA biosynthetic mutants, which is a potential signal to trigger endoreplication in mutant leaves. Present evidence indicates that normal GABA metabolism inhibited endoreplication to prevent immature cell differentiation in leaf development. This research provides a deeper understanding of the role of GABA in plant development.

## 6 Acknowledgments

We apologize if we inadvertently omitted citations of major contributions to this area. We thank Prof. Barry Shelp (University of Guelph, Canada) for generously providing the mutants seeds. We would like to thank Editage (www.editage.cn) for English language editing.

## 7 Author Contributions

YXG studied the root phenotype in relation to GABA metabolism and wrote the draft of the manuscript. HY and YX studied the role of ROS in cell-cycle regulation. GHY studied the role of small molecules in plant development and revised the manuscript.

## 8 Conflict of Interest

The authors declare that the research was conducted in the absence of any commercial or financial relationships that could be construed as a potential conflict of interest.

## 9 Funding

This work was supported by the Special Fund for Basic Scientific Research of Central Colleges, South-Central University for Nationalities (CZP17051), National Natural Science Foundation of China (31270361), studies on the regulation mechanisms of agronomic traits in important crops and *A. thaliana* supported by State Administration of Foreign Experts Affairs in Ministry of Science and Technology of the People’s Republic of China (P193009007) and Fund for Key Laboratory Construction of Hubei Province (Grant No.2018BFC360).

## 10 Data Availability Statement

No datasets were generated for this study.

## Supporting Materials

**Table S1:**
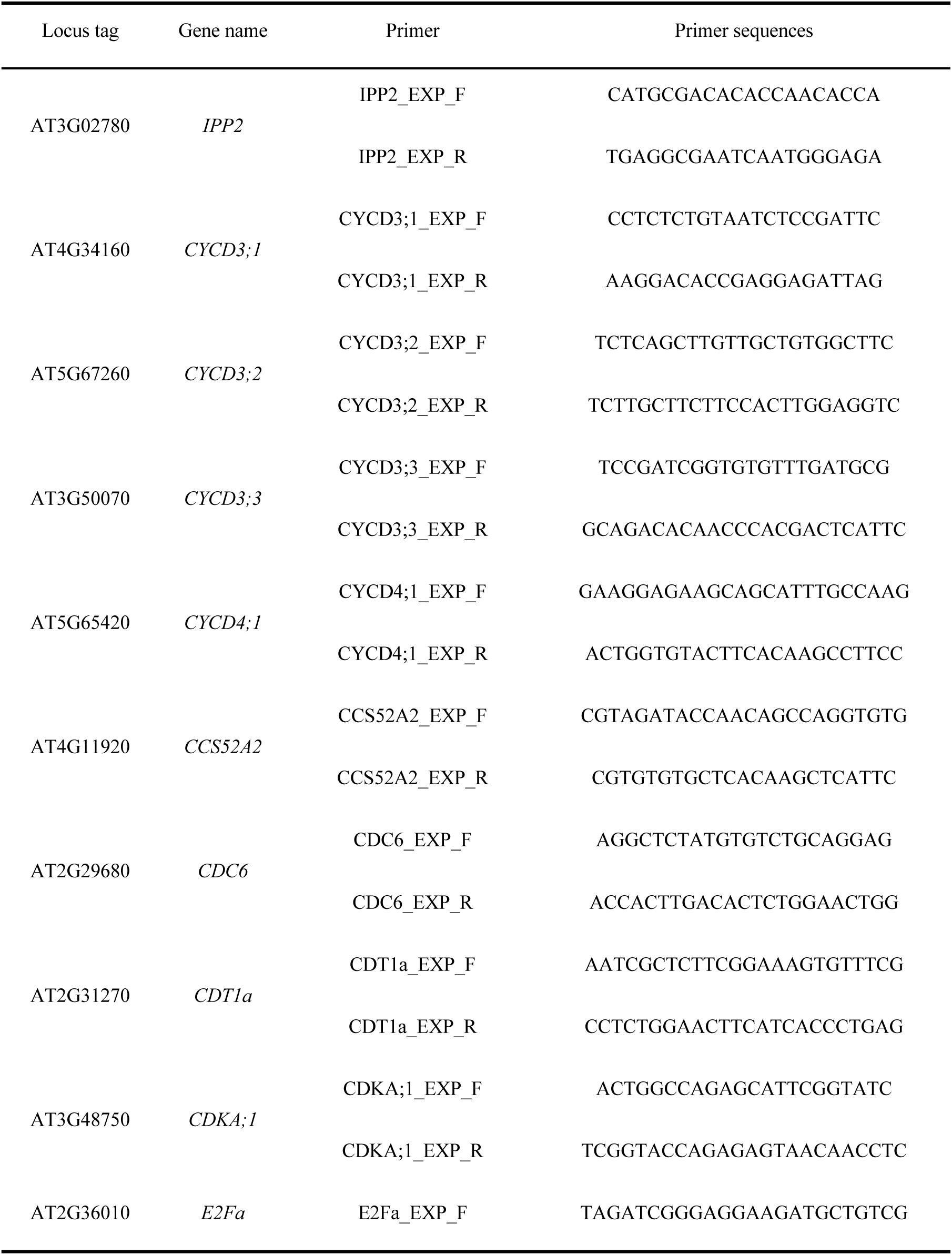
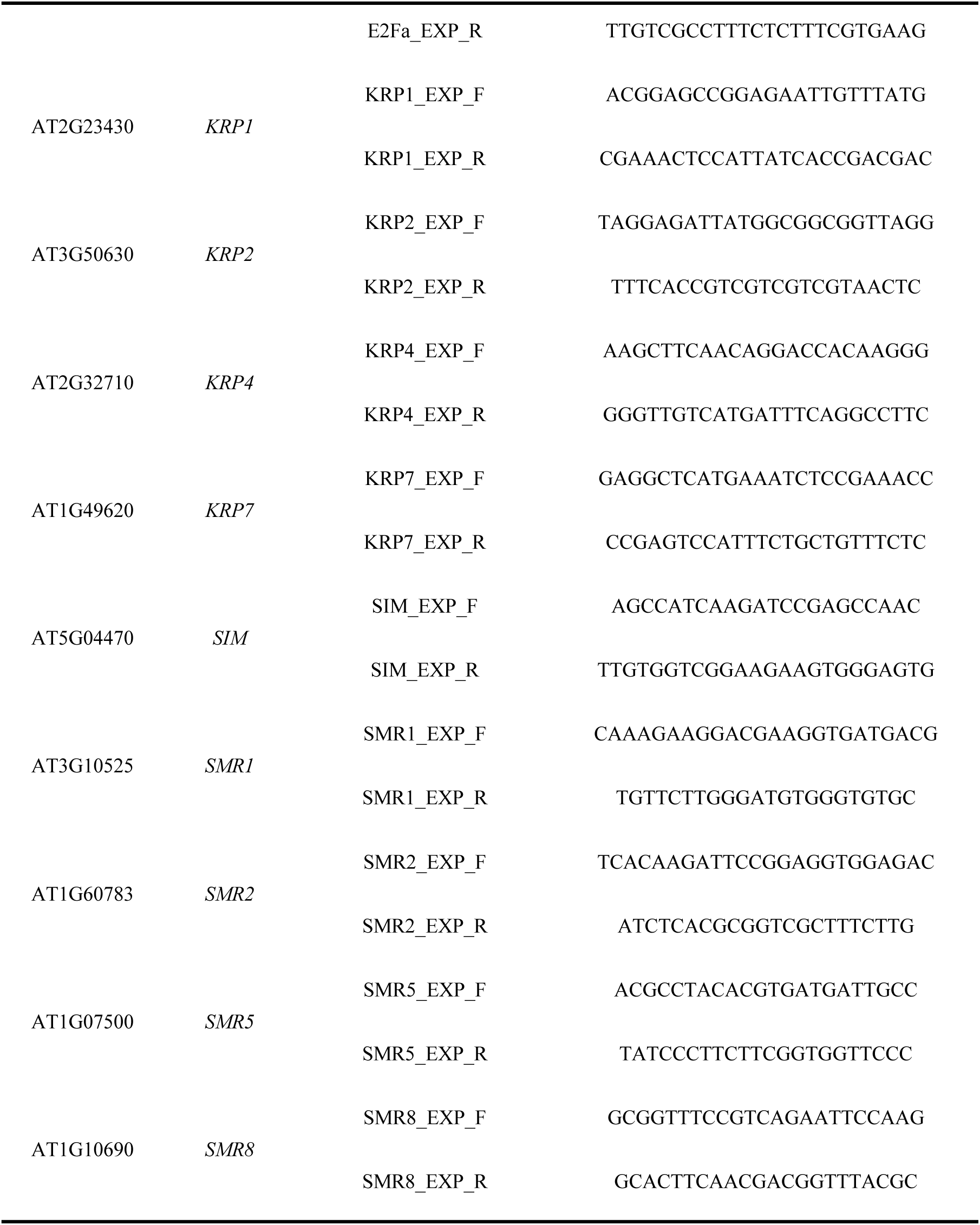
The primers of the genes involved in cell cycle regulation, endocycle initiation, progression and exit.

**Figure S1:**
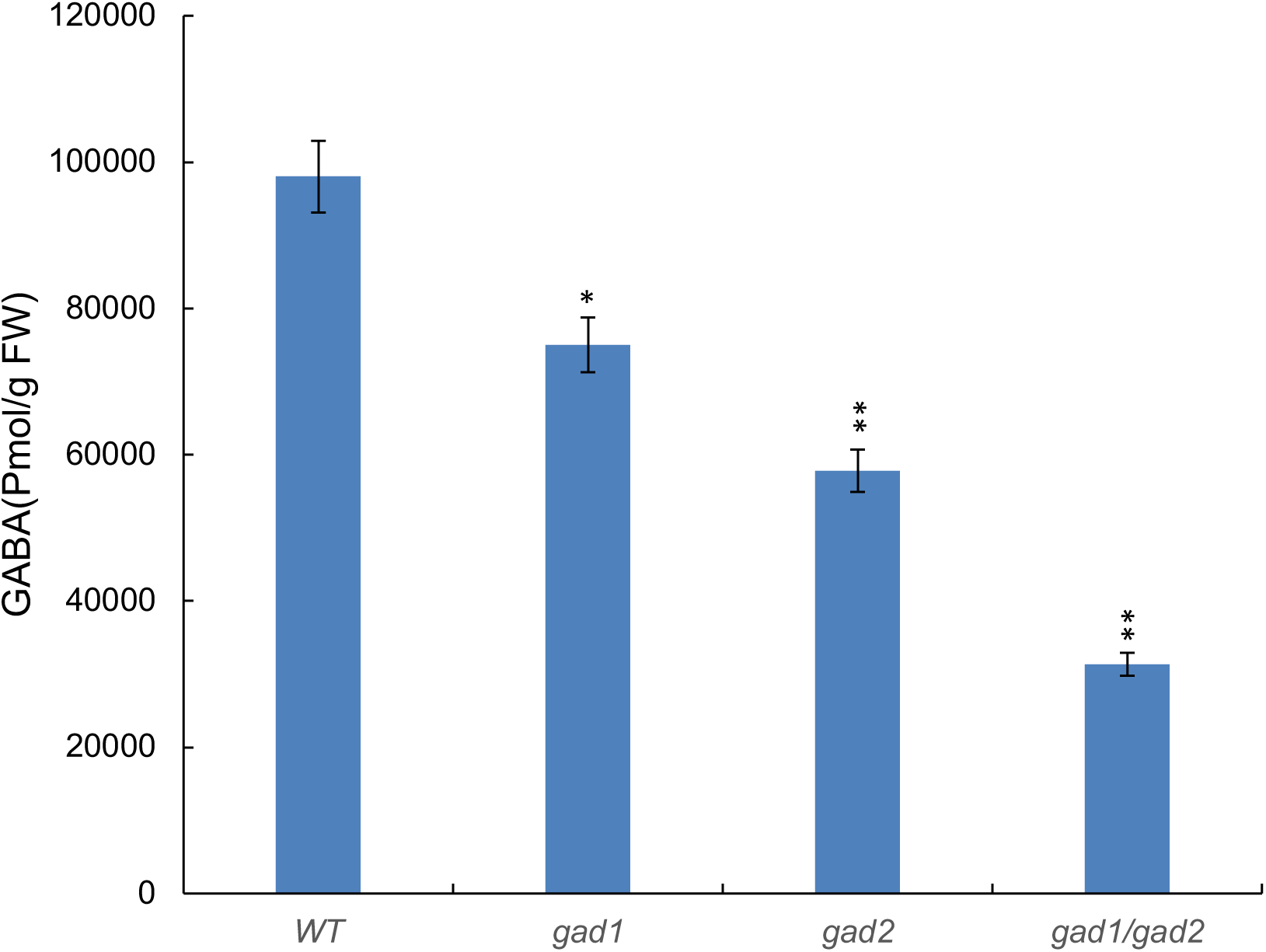
GABA analysis was performed on whole shoot samples. Avarages ± standard error (SE) of three independent replicates. Significant differences from the control are indicated with asterisks: * *P* <0.05, ** *P* < 0.01, by one-way ANOVA.

GABA levels in the leaves of the mutants were determined essentially as described earlier (Allan and Shelp 2006).

## Notes

#### Summary of Updates

Dear Editor, I made correction in the following aspects: 1. In section 7, the author contributions, please replace it by "YXG studied the root phenotype in relation to GABA metabolism and wrote the draft of the manuscript. HY and YXstudied the role of ROS in cell-cycle regulation. GHY studied the role of small molecules in plant development and revised the manuscript." 2. I replaced the reprint online, Thanks for your consideration, Yours, Guanghui

